# Extracellular citrate modulates glutamine metabolism in human macrophages during infection

**DOI:** 10.64898/2026.05.13.724857

**Authors:** Hanna F. Voß-Willenbockel, Felix Leitner, Sina Wischnewski, Stephanie Ng, Kehinde O. Aina, Kristin Metzdorf, Josef Penninger, Hendrikus Garritsen, R. Verena Taudte, Anna Schurich, Michael Steinert, Thekla Cordes

## Abstract

Citrate is a central metabolite linking tricarboxylic acid (TCA) cycle activity to energy and lipid metabolism and supports the synthesis of inflammatory mediators, including itaconate, in macrophages. While citrate is primarily generated endogenously, extracellular citrate levels are elevated under pathological conditions such as citrate transporter disorder. Cells import extracellular citrate through SLC13 transporters, including the sodium-dependent citrate transporter NaCT (encoded by *SLC13A5*). However, whether macrophages take up extracellular citrate and how this affects metabolism and function remains unclear. Here, we combined mass spectrometry and tracing approaches to investigate the metabolic fate of citrate in human macrophage cell lines, primary, and iPSC-derived macrophages. We demonstrate that cells take up extracellular citrate, which was enhanced under metabolic stress conditions. Exogenous citrate was not substantially utilized as a carbon source but selectively altered glutamine metabolism and responses to bacterial infection with *Salmonella enterica* Typhimurium and *Legionella pneumophila* Corby. Our work identifies extracellular citrate as a context-dependent regulator in macrophages that decouples uptake from metabolic utilization.

**Highlights:**

- Macrophages import extracellular citrate via SLC13 transporters
- Extracellular citrate accumulates under hypoxia and inflammatory activation
- Extracellular citrate does not fuel central carbon metabolism in human macrophages
- Citrate modulates glutamine immunometabolism and modulates immune responses

**eTOC blurb:** Voß-Willenbockel et al. demonstrate that human macrophages accumulate extracellular citrate without using it as a major carbon source. Instead, citrate modulates glutamine utilization, inflammatory responses, and host-pathogen interactions revealing a context-dependent regulatory role for extracellular metabolites in immune cell function.

## Introduction

Extracellular citrate is increasingly recognized as a regulator of metabolic and disease processes. Citrate links tricarboxylic acid (TCA) cycle activity to energy metabolism and lipid metabolism, and contributes to processes such as metal chelation and posttranscriptional modification^1–3^. Most cell types synthesize citrate de novo from acetyl-Coenzyme A (CoA) and oxaloacetate in the TCA cycle or via reductive carboxylation^1,3–5^. In addition, cells import extracellular citrate through plasma membrane transporters of the SLC13 family, including the sodium-dependent dicarboxylate transporters NaDC1 (encoded by *SLC13A2*) and NaDC3 (encoded by *SLC13A3*), and the sodium-dependent citrate transporter NaCT (encoded by *SLC13A5*). Citrate is the preferred substrate of the NaCT, which is highly expressed in the liver and brain^1,3,6–9^.

Impaired citrate transport has metabolic and functional consequences. *SLC13A5* impairment in the liver is beneficial in the context of diet-induced obesity or metabolic dysfunction-associated steatotic liver disease (MASLD, formerly NAFLD)^10,11^. In contrast, SLC13A5 transporter disorder with impaired NaCT activity in the brain is a rare metabolic disease associated with delayed brain development and epileptic seizures^1,3,12^. In addition, impaired NaCT activity is associated with increased citrate concentrations in the plasma and cerebrospinal fluid (CSF)^13,14^. However, the impact of elevated extracellular citrate on tissue metabolism, cellular function, and metabolic homeostasis remains poorly understood.

In the tumor microenvironment, high citrate concentrations and malfunction of NaCT reduce tumor growth by altering glycolysis^15,16^. Moreover, extracellular citrate induces cytokine secretion in mice and macrophage cell models, consistent with its role as a damage-associated molecular pattern (DAMP)^16–18^. However, it remains unclear whether macrophages take up extracellular citrate and whether it is used as a carbon source or instead acts as a regulator of cellular metabolism and function. Most commonly used cell culture media lack citrate, suggesting that its contribution to in vitro macrophage metabolism may be underestimated in cultured models^8^. In contrast, citrate is abundant in plasma with 150 - 200 µM^19^ indicating that extracellular citrate may exert systemic metabolic effects that are not captured in standard in vitro models and may influence immune cell function.

In this study, we combined gas chromatography coupled to mass spectrometry (GC-MS) with stable isotope labeling and infection approaches to investigate the impact of extracellular citrate on macrophage metabolism and host-pathogen interactions. We quantified citrate uptake and expression of plasma membrane citrate transporters in macrophages under metabolic stress conditions, including hypoxia and pro-inflammatory stimulation. In addition, we deciphered host-pathogen interactions using intracellular infection models with *Legionella pneumophila* Corby (*L. pneumophila*) and *Salmonella enterica* Typhimurium (*S.* Typhimurium). Together, our study demonstrates the role of extracellular citrate in macrophage metabolism under inflammatory, hypoxic, and infectious conditions. The results may also help to identify new targets for future treatments of diseases associated with altered citrate metabolism and transport, including hepatocellular cancer and brain disorders.

## Materials and Methods

### THP-1 cell cultures

Human monocytic THP-1 cells (ATCC, TIB-202) were cultured in Roswell Park Memorial Institute (RPMI 1640) medium (Gibco, Cat.#21875-034) supplemented with 10 % heat-inactivated fetal bovine serum (FBS) (Bio&Cel, sourced from South America, FBS.SAM.0500, LOT: BS.338199D) and 50 U/ml penicillin-streptomycin (P/S) (Gibco, Cat.#15140-122, Lot: 225194). All cells were cultured at 37 °C with 5 % CO_2_ and 21 % O_2_ in a humidified cell incubator (Thermo Scientific, Heracell Vios 160i CO_2_). THP-1 cells were seeded at a density of 500,000 cells/well, and human monocyte-derived macrophages (hMDMs) with 1 million cells/well in RPMI medium on 12-well tissue culture plates. THP-1 cells were differentiated to macrophage-like cells using 100 ng/ml phorbol 12-myristate 13-acetate (PMA) (BioGems, Cat.#1652981) for 48 h in RPMI medium, rinsed, and cultured for an additional 24 h in RPMI medium before starting treatments. All media were adjusted to pH 7.2. Cells were tested negative for mycoplasma using MycoAlert Mycoplasma Detection Kit (Lonza, Cat.#LT07-118).

### Primary human peripheral blood mononuclear cell (PBMC) cultures

hMDMs were isolated from whole human blood taken from random donors provided by the donation service at the Klinikum Braunschweig. For isolation of PBMCs, whole blood was rinsed out of the leukoreduction system (LRS) chamber with 1x phosphate-buffered saline (PBS) (Gibco, Cat.#18912-014, Lot: 3068235) into 50 ml reaction tubes. For initial separation, blood/PBS solution was layered onto 4 ml Biocoll separation solution (Bio&Sell, BS.L 6115, Lot: BS.9285180) to perform a density gradient centrifugation at 700 *x g* without break. The white PBMC ring was taken off and washed twice with MACS buffer (autoMACS® Rinsing Solution + 5 % MACS® BSA Stock Solution) (Miltenyi Biotec, rinsing solution Cat.#130-091-222, Lot: 7211200253, BSA Cat.#130-091-376, Lot: 7211000566). For further separation, the cells were magnetically labeled with CD14 microbeads (Miltenyi Biotec, Cat.#130-050-201, Lot: 5220200490) in MACS buffer with a 1:5 ratio of cells to CD14 beads and incubated for 15 min on ice. For the separation of CD14-labeled monocytes, MACS LS columns (Miltenyi Biotec, Cat.#130-042-401) attached to the MACS separator (Miltenyi Biotec, MACS®MultiStand, Cat.#130-042-303) were used. Columns were equilibrated with MACS buffer, and CD14-labeled monocytes were eluted with 3 ml MACS buffer using a plunger. For differentiation into macrophages, monocytes were resuspended in RPMI medium containing 10 % FBS, 50 U/ml P/S, 50 U/ml human recombinant colony-stimulating factor (rh M-CSF, Immunotools, Cat.#11343115) and seeded in 6-well tissue culture plates with 2 million cells/ml in 5 ml total volume per well. The cells were incubated for 5 days at 37 °C with 5 % CO_2_ and 21 % O_2_ in a humidified incubator. Differentiated hMDMs were rested for 24 h in RPMI medium before treatment.

### Induced pluripotent stem cell (iPSC) cultures

Human iPSC lines (Healthy Control Human iPSC Line, Female, SCTi003-A; Cat.#200-0511 & Healthy Control Human iPSC Line, Male, SCTi004-A; Cat.#200-0769) were provided by the group for Innovative Organoid Research at the Helmholtz Center for Infection Research, Brunswick. iPSCs were cultured in 6-well plates in mTeSR™ Plus (STEMCELL technologies, Cat.#85850) and medium was changed daily. Cells were washed with 1 ml PBS, treated with 1 ml RELESR-solution (STEMCELL technologies, Cat.#100-0484), and incubated for 2-5 min at room temperature (RT), while observing the cells. Once colonies started to detach, RELESR was removed, and cells were rinsed with 1 ml mTeSR™ until detachment. Destination wells were coated with 1 ml/well Matrigel with growth factors and phenol red (Corning, Cat.#354277) at 37°C for at least 1 h and max. 24 h. Matrigel was removed before adding 1 ml mTeSR™ Plus, and cells were seeded at a split ratio of 1:60 or 1:80. To generate iPSC-derived monocytes, differentiation was performed in multiple stages. Cells were first cultured from day 0 to day 2 in 2 ml/well mesoderm induction medium consisting of IF9S supplemented with 25 ng/ml BMP4, 15 ng/ml Activin-A, and 1.5 µM CHIR 99021. Cells were then cultured from day 2 to day 5 in 3 ml/well hemogenic endothelium induction medium consisting of IF9S supplemented with 50 ng/ml VEGF, 50 ng/ml FGF-2, 10 µM SB43152, and 50 ng/ml SCF. This was followed by culture from day 5 to day 9 in 2 ml/well hematopoietic induction medium consisting of IF9S supplemented with 50 ng/ml VEGF, 50 ng/ml FGF-2, 50 ng/ml SCF, 10 ng/ml IL-3, 50 ng/ml IL-6, and 50 ng/ml TPO. Hematopoietic induction medium was refreshed on day 7. For monocyte induction, the HPCs were gently detached from the well surface by flushing the colonies with monocyte induction medium consisting of IF9S supplemented with 10 ng/ml IL-3, 50 ng/ml IL-6, and 80 ng/ml M-CSF. Floating cells were collected and each well was washed with 1 ml PBS. Subsequently, 0.5 ml TrypLE was added to each well of the six-well plate, followed by incubation for 5 min at 37 °C. Afterward, 1 ml IF9S was added to each well, and cells were detached using a cell scraper and collected in the same tube. An additional 1 ml of IF9S was added to each well, and the remaining cells were collected by gentle pipetting 3-4 times. The pooled cell suspension was centrifuged for 3 min at 300 *x g* at RT. The supernatant was discarded, and the cell pellet was resuspended in 12 ml monocyte induction medium. Cells were seeded at 1 ml/well into a 12-well ultra-low attachment plate and cultured further, with medium changes on differentiation days 12 and 14.

For isolation of iPSC-derived monocytes, wells were washed twice with 1 ml PBS. Cells were then incubated with 1 ml Accutase for 5 min at RT. The reaction was stopped by adding 1 ml Iso-buffer (self-prepared; 1× PBS supplemented with 2 % FBS, 25 mM HEPES, and 2 mM EDTA). Cells were resuspended and transferred into a 15 ml tube and centrifuged for 5 min at 150 *x g*. The cell pellet was then resuspended in 1 ml Iso-buffer and after counting centrifuged again for 5 min at 150 *x g*. Cells were then resuspended in 60 µl Iso-buffer, mixed with 40 µl CD14 micro-beads (Miltenyi Biotec, Cat.#130-050-201), and incubated for 15 min at 4 °C. After incubation, 1 ml Iso-buffer was added, and cells were centrifuged for 5 min at 150 *x g*. The cell pellet was resuspended in 1 ml Iso-buffer and passed through a 100 µm cell strainer. CD14-positive cells were isolated using LS columns placed in a MACS separator. Columns were equilibrated with 3 ml Iso-buffer before loading the cell suspension. After sample application, columns were washed three times with 3 ml Iso-buffer. The columns were then removed from the MACS separator, placed onto new 15 ml tubes, and CD14-positive cells were eluted with 5 ml Iso-buffer using the supplied plunger. After centrifugation for 5 min at 150 × *g*, the cell pellet was resuspended in 1 ml IF9S without growth factors, and cells were counted again. Isolated iPSC-derived monocytes were either used directly for further differentiation or cryopreserved in Bambanker freezing medium (Bambanker, Cat.#BB01). For macrophage differentiation, isolated iPSC-derived monocytes were transferred into 6-well plates and cultured in IF9S supplemented with 80 ng/ml M-CSF. This M-CSF-dependent culture step was used to differentiate iPSC-derived monocytes into iPSC-derived macrophages. iPSC macrophages were treated in Dulbecco’s Modified Eagle Medium (DMEM) with 10 % FBS and 0.5 % P/S.

### Cell treatments

Cells were treated for 24 h with 100 ng/ml lipopolysaccharide (LPS) (Sigma Aldrich, Cat.#L2755, *E. coli* O128:B12, stock 1 mg/ml) and 400 U/ml interferon-gamma (IFNγ) (PeproTech, Cat.#AF-300-02, Lot: 0215AFC27) at 37 °C with 5 % CO_2_ and either 21 % O_2_ (normoxic conditions) or 2 % O_2_ (hypoxic conditions). Cells were treated with 10 μM bis-2-(5-phenylacetamido-1,2,4-thiadiazol-2-yl)ethyl sulfide (BPTES) (CAS.#314045-39-1) or different citrate (Roth, Cat.#X863.1) concentrations as indicated in each figure legend.

### Isotopic tracing analysis

Isotopic tracing analysis of THP-1 and hMDMs was performed in SILAC RPMI 1640 FLEX Medium (Gibco, Cat.#A24942-01) supplemented with 1.14 mM L-Arginine and 0.218 mM L-Lysine hydrochloride. For ^13^C glucose tracing, the medium was supplemented with 11.1 mM [U-^13^C_6_]glucose (Cambridge Isotope Laboratories, Cat.#CLM-1396-1) and 2 mM unlabeled ^12^C glutamine. For ^13^C glutamine tracing, the medium was supplemented with 11.1 mM unlabeled ^12^C glucose and 2 mM [U-^13^C_5_]glutamine (Cambridge Isotope Laboratories, Cat.#CLM-1822-H-0.5). Citrate tracing was performed by supplementing 0.5 mM [2,4-^13^C_2_]citrate (Cambridge Isotope Labs, Cat.#CLM-148) to RPMI medium. All tracer media were sterile-filtered through 0.22 µm filters and supplemented with 10 % FBS, 50 U/ml P/S. Labeling on metabolites from ^13^C tracer was quantified using a GC-MS platform.

### Metabolite extraction for GC-MS analysis

Metabolites were extracted as previously described in detail^20^. Briefly, cells were washed with saline solution (0.9 % w/v NaCl) and quenched with 0.25 ml −20 °C methanol. After adding 0.1 ml 4 °C cold water containing an internal standard norvaline (5 µg/ml). The mixture was collected into a pre-cooled microcentrifuge tube, and 20 µl of the methanol-water mixture was transferred into a 96-well plate for protein quantification. Next, 0.25 ml −20 °C chloroform was added to the sample, and the extracts were vortexed for 10 min at 4 °C and centrifuged at 17,000 × *g* for 5 min at 4°C. 170 μl of the upper aqueous phase was evaporated under vacuum at 4 °C, and the organic phase was dried at room temperature and used for GC-MS analysis. Medium samples were centrifuged at 300 x *g* for 5 min at 4°C, then 10 µl of medium was mixed with 90 µl of extraction fluid (methanol:water is 8:1, containing 4.44 µg/ml norvaline), vortexed for 15 sec, and spun again at 17,000 x *g* for 5 min at 4°C. 80 µl of the supernatant was dried under vacuum at 4°C and used for GC-MS analysis.

### Gas Chromatography – Mass Spectrometry (GC-MS) and sample preparation

Metabolites were analyzed and quantified, as previously described in detail by Cordes and Metallo^20^. Polar metabolites were derivatized by an automated sample robot (MPS2-XL-Twister, Gerstel) with 15 µl 2 % methoxamine hydrochloride (MeOX) (Sigma Aldrich, Cat.#593-56-6) for 90 min at 55 °C with shaking and 15 µl *N*-tert-butyldimethylsilyl-*N*-methyltrifluoroacetamide (MTBSTFA) with 1 % tert-butyldimethylchlorosilane (RESTEK, Cat.#35601) for 60 min at 55 °C with shaking. 1 µl of each sample was injected into a GC-MS equipped with a ZB-35MS UI column (Phenomenex, Cat.#7HG-G003-11-GGA-C). Helium was used as a carrier gas at 1 ml/min rate. The GC oven was held at 100 °C for 2 min. Then, the temperature was increased to 325 °C at 10 °C/min and held for 4 min, followed by 300 °C for an additional 5 min. The MS was operated in electron impact ionization mode at 70 eV. During measurements, the source was held at 230 °C and the quadrupole at 150 °C. The total measurement took 28.5 min, and metabolites were measured in full scan (100 m/z – 650 m/z).

The raw data from GC-MS were analyzed with the help of OpenChrom software (Lablicate) and an in-house integration code using MATLAB 2022b (MathWorks) to compute mass isotopomer distributions and total metabolite abundance by integrating mass fragments with corrections for natural isotope abundances as described in Cordes and Metallo^20^. Metabolite abundances were normalized to internal standard norvaline and protein content per well, as indicated in each figure legend.

### Transesterification and GC-MS measurement of fatty acid methyl esters (FAMEs)

Transesterification of fatty acids to fatty acid methyl esters (FAMEs) was performed as described in Cordes and Metallo^20^. Briefly, 500 µl 2 % v/v H_2_SO_4_ in methanol was added to the dried organic phases for transesterification. The samples were incubated for 2 h at 50 °C before adding 100 µl saturated NaCl and 500 µl hexane. The samples were vortexed and left to sit for 1 min at room temperature for phase separation. The upper hexane phase was transferred into a new 1.5 ml reaction tube. 500 μl hexane was added again for a second extraction round. The hexane phases were dried overnight, resuspended with 75 µl hexane, and transferred into glass vials with inserts. For the measurement of FAMEs, the GC oven was held at 100 °C for 3 min, increased up to 205 °C at 25 °C/min, and then to a temperature of 230 °C at 5 °C/min. The oven temperature was then increased to 300 °C at 25 °C/min and held at that temperature for 2 min. Total measurement time was 17 min.

### Extracellular glucose and lactate measurements

Medium glucose and lactate concentrations were determined using a Yellow Springs Instrument (YSI Inc., YSI 2950). Spent media were centrifuged at 300 x *g,* and the supernatant was again centrifuged at 17,000 x *g* for 10 min to remove impurities. 200 µl of medium sample was analyzed for glucose and lactate, and concentrations were calculated from a linear calibration curve from measured standards.

### Liquid chromatography mass spectrometry (LC-MS) sample preparation and analysis

Metabolite samples for liquid chromatograph mass spectrometry (LC-MS) analysis were extracted with 250 µl −20 °C MeOH containing internal standard (3.1 ng/ml DL-2-fluorophenylglycine and 3.1 ng/ml D_6_-Cholesterol) and 100 µl 4 °C Milli-Q water. Cells were washed before with 1 ml 0.9 % NaCl. The cells were scraped from the plate, vortexed, and 20 µl of the cell suspension was transferred into a 96-well plate for quantifying protein content before 250 µl −20 °C chloroform was added. The samples were inverted three times and vortexed for 10 min. 180 µl of the upper aqueous phase was transferred into a 1.5 ml tube and dried overnight in a speed vacuum concentrator at 4°C.

Immediately before analysis, the samples were reconstituted in 200 µl of the starting eluent and transferred into HPLC vials. Samples were measured on a 1290 Infinity UHPLC system (Agilent Technologies, Waldbronn, Germany) consisting of a binary pump, an autosampler, and a column oven, coupled to a QTRAP 5500 mass spectrometry system (AB SCIEX, Foster City, CA, USA) and controlled using Analyst 1.7 software (AB SCIEX, Foster City, CA, USA). For the separation of metabolites, a HILIC-Z column (Agilent InfinityLab Poroshell 120, 2.1 × 150 mm, 2.7 μm) with a guard column (Agilent InfinityLab Poroshell 120, 2.1 × 5 mm, 2.7 μm) was used. The column temperature was kept at 25 °C, whereas the autosampler temperature was 4 °C. Injection volume was 5 µl. The gradient consisted of eluent A (10 mM HCOONH_4_ in H_2_O (0.1 % FA), pH 3.5) and eluent B (10 mM HCOONH_4_ in 1:9 H_2_O/ACN (0.1% FA). The mass spectrometer was operated in positive ionization mode. Curtain gas was at 40 psi, collision-activated dissociation gas at medium, and the voltage at + 4500 V. Temperature was at 450 °C, and GS1 and GS2 at 45.5 and 40.0 psi, respectively. The scan rate was 200 Da/s. Metabolites were identified using optimized MRM conditions for each compound. For quantification, methods were created using MultiQuant 3.0 software (AB SCIEX, Foster City, CA, USA) containing the retention times of the metabolites from the LC method and enabling automatic integration.

### Respirometry

Respiration was measured in adherent monolayers of cells using an Agilent Seahorse XF96 Analyzer and the Seahorse XFe96/XF Pro FluxPak (Agilent Technologies, Cat.#103792-100). Cells were seeded at a density of 300,000 to 400,000 cells/well with cellular replicates per condition and assayed in RPMI 1640 medium (Sigma Aldrich, Cat.#R1383) supplemented with 2 mM HEPES, 8 mM glucose, 2 mM glutamine, and 2 mM sodium pyruvate. Cells were washed twice with 100 µl of assay medium, and 150 µl of assay medium was added to each culture well. Respiration was measured under basal conditions as well as after injection of 2 µM oligomycin, two sequential additions of 1 µM trifluoromethoxy carbonyl cyanide phenylhydrazone (FCCP), and addition of 0.5 µM rotenone and 1 µM antimycin A (from Streptomyces, Sigma-Aldrich, Cat.#A8675-50MG). The assay medium contained different citrate concentrations, and pH was adjusted to 7.2 using NaOH. The raw data were analyzed by Wave 2.4.3 software and data were normalized to the protein content of each well.

### Quantitative real-time polymerase chain reaction (qRT-PCR)

RNA isolation was performed according to the manufacturer’s instructions for the NucleoSpin^®^ RNA kit (Macherey-Nagel, Cat.#740955.250). The RNA concentration was quantified by using a microplate reader (Tecan Infinite^®^ 200 Spark^®^). 10 µl RNA (100 ng/µl) per sample was reverse transcribed into cDNA using High-Capacity cDNA Reverse Transcription kit (Thermo Fisher Scientific, Cat.#4368813). The reaction was done in a Thermocycler (Professional, Biometra®), holding the temperature at 25 °C for 10 min, then increasing it to 37 °C for 120 min before finally heating to 85 °C and holding the temperature for 5 min. Individual 10 μl SYBR Green real-time PCR reactions consisted of 2 μl of diluted cDNA (50 ng), 5 μl iTaq Universal SYBR Green Supermix (BIO-RAD, Cat.#1725124), and 1.5 μl of each 10 μM forward and reverse primers. PCR was carried out in 96-well plates (MicroAmp™ Optical 96-Well Reaction Plate, Applied Biosystems by Life technologies^®^, Cat.#N8010560) and centrifuged at 300 x *g* for 30 sec at RT before measurement with QuantStudio Design & Analysis Software v1.5.2 (Thermo Fisher Scientific) using a three-stage program provided by the manufacturer: 95 °C for 3 min, 45 cycles of 95 °C for 10 sec, and 60 °C for 30 sec. The cycling threshold (CT) values were normalized to the average CT value of *RPL27* as a housekeeping gene, and the fold change of the ΔΔCT was calculated.

Primer uses are as follows. *RPL27* forward: TGGACAAAACTGTCGTCAATAAGG; *RPL27* reverse: AGAACCACTTGTTCTTGCCTGTC; *SLC13A3* forward: CCATTGAGGAGTGGAACCTGCA; *SLC13A3* reverse: GTGTTGCTCAGCCACATGGACA; *SLC13A5* forward: CTGCCACTCGTCATTCTGATG; *SLC13A5* reverse: ATGTTGGTGTCCTTCATGTACTG, *SLC7A5* forward: GGAAGGGTGATGTGTCCAATC; *SLC7A5* reverse: TAATGCCAGCACAATGTTCCC.

### Enzyme-linked immunosorbent assay (ELISA)

The enzyme-linked immunosorbent assay (ELISA) was performed following the manufacturer’s instructions of ELISA MAX^TM^ Deluxe set for human Interleukin 6 (IL-6) (BioLegend, Cat.#430504). Medium was diluted 1:150 in assay diluent A, and samples were measured in duplicates. The horseradish peroxidase (HRP) reaction was stopped by adding 2 N H_2_SO_4_ before absorbance measurement at 450 nm and 570 nm in a Tecan Infinite® 200 Spark plate reader. Data was normalized by subtracting the absorbance at 570 nm from the absorbance at 450 nm.

### Immune markers on T-cells

Cryopreserved PBMCs were thawed and cultured at 1×10⁶ cells/ml in complete RPMI (cRPMI) medium (ThermoFisher, 31870-025, 10 % FBS, 1 % glutamine, 0.1 mM non-essential amino acids, 10 mM HEPES buffer, and 1 mM sodium pyruvate) supplemented with either 0 mM, 0.2 mM, 1 mM, or 5 mM citrate (Sigma-Aldrich, Cat.#C0759) and 20 IU/ml IL-2 (Proleukin). After 24 hours, cells were left unstimulated or activated in plates coated with anti-CD3 (BioLegend, Cat.#344818) and soluble anti-CD28 (BioLegend, Cat.#302934) antibodies for 24 hours. For T-cell phenotyping, brefeldin-A (BioLegend, Cat.#420601) was added to a final concentration of 1 µg/ml, and cells were incubated for the final 15 hours at 37 °C with 5 % CO₂. Cells were then washed with 1x PBS and stained for surface markers using the antibodies, anti-CD3 (BioLegend, Cat.#344818) and anti-CD62L (BioLegend, Cat.#304843) as well as with blue viability dye (ThermoFisher, Cat.#L23105) in 1x PBS for 30 minutes at 4°C in the dark. Cells were washed with 1x PBS and fixed with 4 % paraformaldehyde (Severn Biotec Ltd, Cat.#40-7401-05) for 20 minutes at room temperature in the dark. Following fixation, cells were washed with 1x PBS and permeabilized in 0.1 % saponin buffer for 15 minutes at room temperature before staining for intracellular markers using the antibody, anti-TNF-α (Cat.#50913) in saponin buffer for 30 minutes at 4 °C in the dark. After a final wash, cells were resuspended in PBS before acquisition. Samples were acquired on a BD Fortessa and data were analysed using FlowJo v10 software.

### Bicinchoninic protein assay (BCA)

Protein concentrations were quantified using the Pierce bicinchoninic protein assay (BCA) kit (Thermo Scientific, Cat.#23250) following the manufacturer’s instructions. Briefly, dried samples from the metabolite extraction in the 96-well cell culture plate were dissolved in 1x radioimmunoprecipitation assay (RIPA) buffer. The BCA assay for the respirometry assay was performed on the respirometry cell plate after the medium was removed and the cells were dissolved in 25 µl 1x RIPA buffer.

### Co-culture infection experiments

Infection experiments with *Salmonella enterica* Typhimurium (*S.* Typhimurium) (ATCC 19585) and *Legionella pneumophila* Corby (*L. pneumophila*)^21^ were performed with differentiated macrophage-like THP-1 cells in a humidified incubator at 37 °C with 5 % CO_2_ using RPMI medium without P/S supplementation. *L. pneumophila* wild-type (WT) and the non-infectious *ΔdotA* mutant were grown for 72 h on buffered charcoal yeast extract-agar (BCYE) (10 g ACES (N-(2-acetamido)-2-aminoethanesulfonic acid), 10 g yeast extract, 0.4 g L-cysteine, 0.25 g iron (III) nitrate, and 15 g agar in 1000 ml ddH_2_O)^22^ at 37 °C. Differentiated THP-1 cells were infected with *L. pneumophila* at a multiplicity of infection (MOI of 1:10) and incubated for 2 h. Cells were washed with 1x PBS and further cultured in tracer medium for 22 h. For infection with *S.* Typhimurium, an overnight culture grown on lysogen broth (LB) agar plates was used. Differentiated THP-1 cells were infected at an MOI of 1:1, cultured for 1 h, washed with 1x PBS, and further incubated in 750 µl RPMI medium with 100 µg/ml gentamicin (Gibco, Cat.#15710-064) without P/S for 1 h. To quantify colony-forming units (CFU), THP-1 cells were lysed using 1 % triton x100 (alkylphenylpolyethylenglykol) (Fluka Chemie AG, Cat.#274173 687) for *L. pneumophila* infection or sterile water for *S.* Typhimurium infection for 2-3 min. 20 µl of bacterial suspension with serial dilutions of 10^0^, 10^−1^, 10^−2^, and 10^−2^ were used for inoculation and CFU analysis. The *L. pneumophila* dilutions were cultivated on BCYE-agar for 72 h at 37 °C. The *S.* Typhimurium dilutions were cultivated on LB-agar overnight at 37 °C. CFU was calculated from counted colonies.

### Bacterial growth curves

Bacterial growth was quantified in 96-well plates with 200 µl bacterial suspension. For *S.* Typhimurium analysis, an overnight culture was prepared in LB medium and used to inoculate a pre-culture in RPMI 1640 medium with 10 % FBS. The growth curve was determined in RPMI 1640 medium, with 10 % FBS and a starting OD_600_ of 0.1. Cultures were supplemented with 1 mM citrate, 2 mM citrate, or 10 µM BPTES. OD_600_ was measured every 15 min over 24 h, at 37 °C with shaking in between at 182.6 rpm in a plate reader (Tecan Infinite M200). For *L. pneumophila* analysis, an overnight culture was prepared in yeast extract beef (YEB) (37 °C; 180 rpm) (10 g ACES, 10 g yeast extract, 0.4 g L-cysteine, 0.25 g iron (III) pyrophosphate in 1000 ml dd H_2_O) medium, which was used to inoculate the growth culture in YEB. The growth curve was determined with a starting OD_600_ of 0.1. Cultures were supplemented with 0.5 mM or 1 mM citrate. OD_600_ was measured every 60 min over 68 h, at 37 °C with shaking in between at 182.6 rpm in a plate reader (Tecan Infinite M Nano).

### Data analysis and statistics

Data visualization and statistical analysis were performed using GraphPad Prism (v10.3.1, GraphPad Software), Adobe Illustrator CS6 (v24.1.2, Adobe Inc.), and BioRender.com. The type and number of replicates and the statistical test used are described in each figure legend. Data are presented as means ± standard error of mean (s.e.m.). Experiments were independently repeated as indicated in the figure legend. *P* values were calculated using an unpaired *t*-test, one-way ANOVA, or two-way ANOVA with Fisher’s least significant difference (LSD) post hoc test and without correction for multiple comparisons as indicated in each figure legend. Normality of data was tested with Shapiro-Wilk test. For all tests, *p* < 0.05 was considered significant with \**p* < 0.05; \*\**p* < 0.01; \*\*\**p* < 0.001, and ^#^*p* < 0.0001.

## Results

### Human macrophages express SLC13 transporter and accumulate extracellular citrate

Uptake of extracellular citrate is implicated in disease-relevant processes, specifically in liver and brain tissue with high *SLC13A5* expression^15–18,23^. Citrate also plays a central role in macrophage metabolism, for instance, by providing carbons for itaconate synthesis^24^. Most common cell culture media, including RPMI medium, lack citrate, though citrate is abundant in plasma at sub-millimolar concentrations. Thus, extracellular citrate may exert systemic effects on immune responses that are not captured in standard in vitro models^25^. However, whether human macrophages take up citrate from the extracellular environment remains unclear. To address this, we cultured human macrophages in RPMI medium supplemented with extracellular citrate (1 mM) and quantified citrate levels and metabolism using GC-MS. Environmental conditions like hypoxia impact TCA cycle metabolites and lead to decreased citrate abundance in macrophages^26^. Therefore, we hypothesized that citrate uptake is influenced by cellular state. To test this hypothesis, we analyzed citrate uptake under hypoxic conditions (2 % oxygen) and following inflammatory stimulation with lipopolysaccharide and interferon-gamma (LPS/IFNγ) in differentiated macrophage-like THP-1 cells.

We observed that hypoxic conditions decreased intracellular citrate levels in PMA-differentiated macrophage-like THP-1 cells (Fig. 1a). Citrate levels were increased upon citrate supplementation, with a more profound effect under hypoxia. Further, pro-inflammatory stimulation with LPS/IFNγ increased intracellular citrate levels in hypoxia compared to normoxia (Fig. 1a). Consistently, we observed minimal citrate uptake under normoxia, whereas uptake was significantly enhanced under hypoxia (Fig. 1b). These data indicate that citrate uptake is influenced by cellular state and is enhanced under metabolic stress conditions suggesting regulatory mechanisms to maintain metabolic homeostasis.

**Figure 1:**
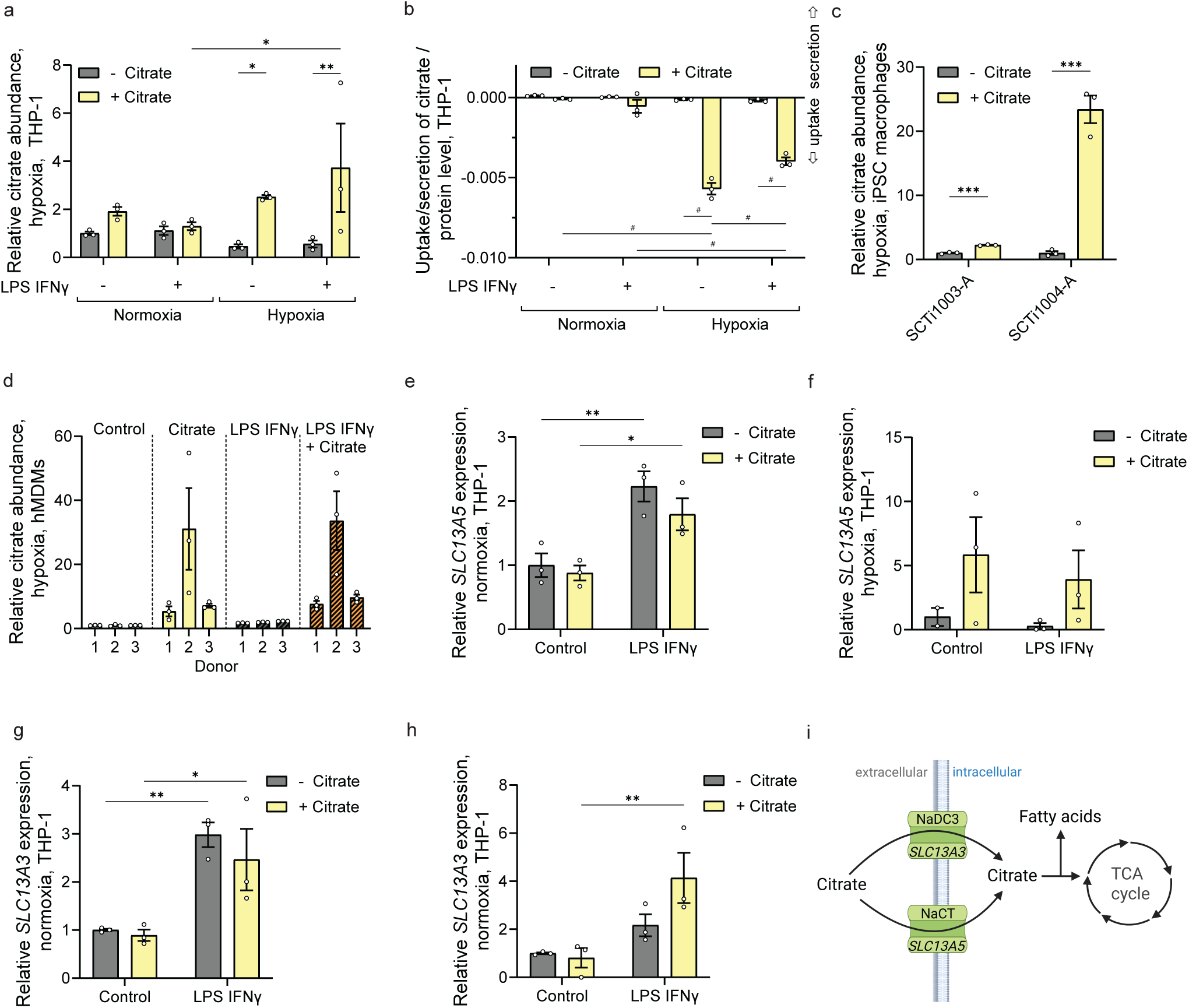
Human macrophages express SLC13 transporter and accumulate extracellular citrate. **a**, Relative intracellular abundance of citrate in macrophage-like THP-1 cells with and without LPS/IFNγ and citrate in normoxic and hypoxic conditions. **b**, Accumulation of extracellular citrate by macrophage-like THP-1 cells with and without LPS/IFNγ and citrate in normoxic and hypoxic conditions. **c**, Relative intracellular abundance of citrate in two strains of iPSC-derived macrophages from two iPSC lines, SCTi1003-A and SCTi1004-A, treated with LPS/IFNγ in hypoxic conditions. **d**, Relative intracellular abundance of citrate in hMDMs from 3 random donors with LPS/IFNγ in hypoxic conditions. **e,f,** Expression of *SLC13A5* in macrophage-like THP-1 cells in **e,** normoxic and **f,** hypoxic conditions. **g,h**, Expression of *SLC13A3* in macrophage-like THP-1 cells in **g,** normoxic and **h,** hypoxic conditions. **i**, Schematic depicting uptake of extracellular citrate via NaCT (*SLC13A5*) and NaDC3 (*SLC13A3*) into the cytosol and potential routes for TCA cycle and fatty acid metabolism. Experiments were independently repeated two (c,e,g) or more times (a,b,f,h) or with 3 donors (d) with each three cellular replicates. Data baseline corrected to control conditions without citrate (a,b-h). RT-qPCR data depicted as ΔΔCT relative to control condition (e-g). Cells were treated with 1 mM citrate and exposed to 100 ng/ml LPS and 400 U/ml IFNγ for 24 hours. Hypoxic conditions were performed with 2 % oxygen. Data are presented as means ± s.e.m.. Statistical significances were determined with two-way *ANOVA* (a,b, e-g) or uncorrected Fisher’s LSD test (c) with * *p* < 0.05; ** *p* < 0.01, *** *p* < 0.001, ^#^ *p* < 0.0001.

Next, we validated these findings with more physiologically relevant models using human iPSC-derived macrophages (Fig. 1c) and human monocyte-derived macrophages (hMDMs) obtained from three healthy donors (Fig. 1d). We observed significantly increased intracellular citrate levels in cultures with supplemented citrate, confirming that citrate accumulation is a conserved feature of human macrophages.

To investigate the molecular basis of citrate uptake, we analyzed the expression of SLC13 family transporters implicated in citrate transport, namely NaDC3 (encoded by *SLC13A3*) and NaCT (encoded by *SLC13A5*)^3^. We found that *SLC13A5* expression increased upon LPS/IFNγ stimulation under normoxic conditions and was further induced by extracellular citrate under hypoxia (Fig. 1e, f). These data indicate that human macrophages express *SLC13A5* and that its expression is regulated by extracellular and inflammatory cues. Furthermore, inflammatory stimulation increased *SLC13A3* expression under both normoxic and hypoxic conditions, whereas extracellular citrate had only minor effects (Fig. 1g, h). These findings suggest that SLC13 transporters may contribute to citrate uptake and accumulation in human macrophages. Further, citrate itself and inflammatory stimulation may influence citrate transporter expression (Fig. 1i).

Since extracellular citrate uptake and transporter expression are regulated by inflammatory stimuli, we next examined the metabolic consequences of extracellular citrate on human macrophages. Under normoxic conditions, citrate supplementation had minimal effects on metabolite levels, including TCA cycle intermediates and amino acids, in both control and LPS/IFNγ-treated cells (Sup. Fig. 1a-c). Citrate is an allosteric inhibitor of glycolysis by targeting 6-phosphofructo-1-kinase (PFK-1)^3,15,16^. We therefore quantified glycolytic activity by measuring glucose uptake and lactate secretion in response to extracellular citrate. We did not observe significant effects on glucose uptake or lactate secretion, indicating that extracellular citrate does not significantly alter glycolysis under these conditions (Sup. Fig. 1d, e). To assess potential effects on oxidative phosphorylation, including potential inhibition of succinate dehydrogenase (SDH), we quantified respiration in real time^27,28^. Citrate did not affect oxygen consumption rate (OCR) of macrophages at 1.25 mM citrate (Sup. Fig. 1f, g), whereas a higher concentration of 5 mM citrate significantly reduced maximal OCR (Sup. Fig. 1h, i). These data indicate that extracellular citrate has minor effects on glycolysis and central carbon metabolism under physiological conditions in our experimental setup. Collectively, these findings establish that extracellular citrate is taken up by human macrophages in a condition-dependent manner, which provides a rationale to investigate its metabolic fate and functional consequences.

### Extracellular citrate does not fuel central carbon metabolism but alters glutamine metabolism in human macrophages

Since macrophages accumulate extracellular citrate with minimal effects on steady-state metabolite levels, we next investigated how citrate influences metabolic fluxes in macrophage metabolism. To quantify the fate of citrate, we cultured human macrophages in the presence of 0.5 mM [2,4-^13^C_2_]citrate under hypoxic conditions, where citrate uptake is enhanced (Fig. 1). The abundance of most TCA cycle intermediates and amino acids was not significantly affected by extracellular citrate (Sup. Fig. 2a, b). If utilized metabolically fueling central carbon metabolism, we expect labeling from citrate on TCA cycle intermediates and fatty acids, including palmitate (Fig. 2a). However, we did not detect significant labeling on TCA cycle intermediates, related amino acids, or palmitate (Fig. 2b). These data indicate that extracellular citrate is not substantially used as a carbon source for TCA cycle metabolism in our cell models.

**Figure 2:**
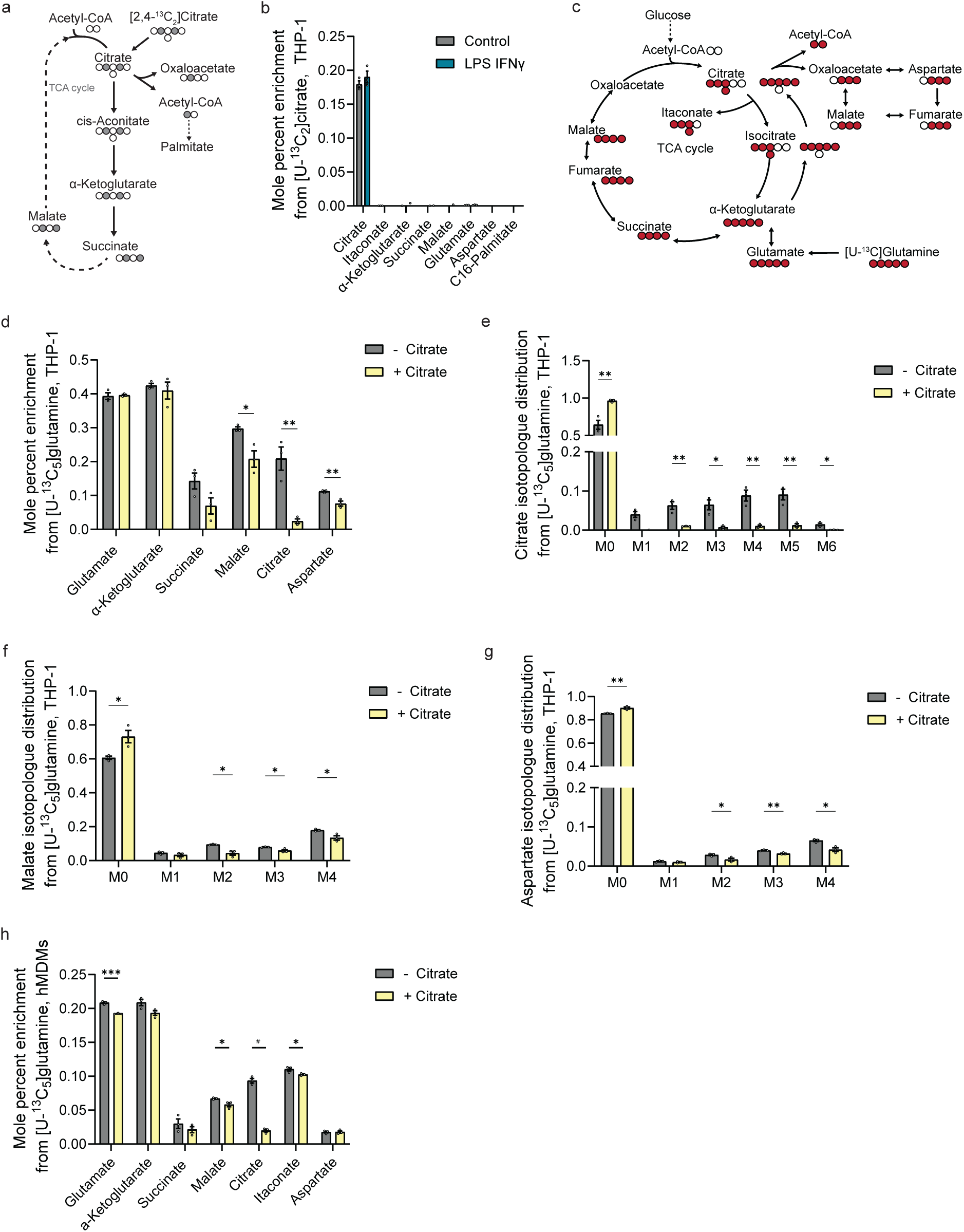
Citrate is not a major carbon source but alters glutamine metabolism in human macrophages under hypoxic conditions. **a**, Schematic depicting atom transition of [2,4-^13^C_2_]citrate in TCA cycle and fatty acid metabolism. Open circles depict ^12^C atoms; grey circles ^13^C atoms**. b**, Mole percent enrichment of TCA cycle intermediates and palmitate from 0.5 mM [2,4-^13^C_2_]citrate. **c,** Schematic depicting atom transition of [U-^13^C_5_]glutamine for TCA cycle metabolism. Open circles: ^12^C atoms; red circles: ^13^C atoms. **d,** Mole percent enrichment from [U-^13^C_5_]glutamine on TCA cycle intermediates with and without extracellular citrate in macrophage-like THP-1 cells. **e-g,** Isotopologue distribution from [U-^13^C_5_]glutamine on **e**, citrate, **f**, malate, and **g**, aspartate. **h,** Mole percent enrichment from [U-^13^C_5_]glutamine on TCA cycle intermediates with and without extracellular citrate in hMDMs. Cells were treated with 1 mM citrate and exposed to 100 ng/ml LPS and 400 U/ml IFNγ for 24 hours. Hypoxic conditions were performed with 2 % oxygen. Experiments were repeated independently one (b) or three (d-h) times with each three cellular replicates. Data are presented as means ± s.e.m.. Statistical significances were determined with uncorrected Fisher’s LSD test with * *p* < 0.05; ** *p* < 0.01, *** *p* < 0.001, ^#^ *p* < 0.0001.

Small molecules such as citrate and itaconate can act as metabolic regulators that influence metabolic fluxes and substrate utilization without altering steady-state metabolite levels or serving as carbon sources. Further, pro-inflammatory conditions drive metabolic reprogramming in macrophages, including an increase in glutamine anaplerosis to fuel increased succinate generation^29^. Therefore, we focused on glutamine-dependent fluxes and applied [U-^13^C_5_]glutamine tracer under hypoxic and pro-inflammatory conditions to quantify incorporation into central carbon metabolism in macrophage-like THP-1 cultures (Fig. 2c). We observed that extracellular citrate decreased incorporation of glutamine-derived carbons into TCA cycle intermediates and related amino acids, specifically on citrate, malate, and aspartate (Fig. 2d). As expected, isotopologue distribution on citrate was significantly decreased due to the dilution of the citrate pool by the applied unlabeled citrate (Fig. 2e). Isotopologue distributions of malate and aspartate were significantly reduced with citrate with comparable decreases on M3 and M4 labeling (Fig. 2f, g). These data suggest that both oxidative (M4) and reductive (M3) parts of the TCA cycle are affected by extracellular citrate. We confirmed reduced ^13^C glutamine utilization in hMDMs (Fig. 2h, Sup. Fig. 2c, d). In contrast to glutamine, our [U-^13^C_6_]glucose tracer studies revealed no change in glucose utilization for TCA cycle metabolism, aside from the expected reduction in citrate labeling (Sup. Fig. 2e, f). Collectively, these results demonstrate that human macrophages accumulate extracellular citrate but rather than using it as a major carbon source, citrate selectively rewires glutamine-dependent TCA cycle metabolism under stress conditions. Furthermore, they demonstrate that metabolic fluxes can be changed without affecting metabolic abundances.

### Extracellular citrate modulates inflammatory responses

Since extracellular citrate reprograms glutamine-dependent metabolism without contributing to central carbon metabolism (Fig. 2), we next asked whether these metabolic changes translate into functional effects of macrophage activation. Metabolic reprogramming is essential for macrophage function^24^, and extracellular citrate influences cytokine production^16–18^. Thus, we quantified interleukin-6 (IL-6) secretion in LPS/IFNγ-stimulated macrophage-like THP-1 cells in the presence or absence of extracellular citrate to get insights into immune cells. Under hypoxic conditions, IL-6 concentrations were significantly increased upon citrate supplementation (Fig. 3a), while no effects were observed in normoxic conditions (Fig. 3b). This finding is consistent with the increased citrate uptake in hypoxic conditions (Fig. 1).

**Figure 3:**
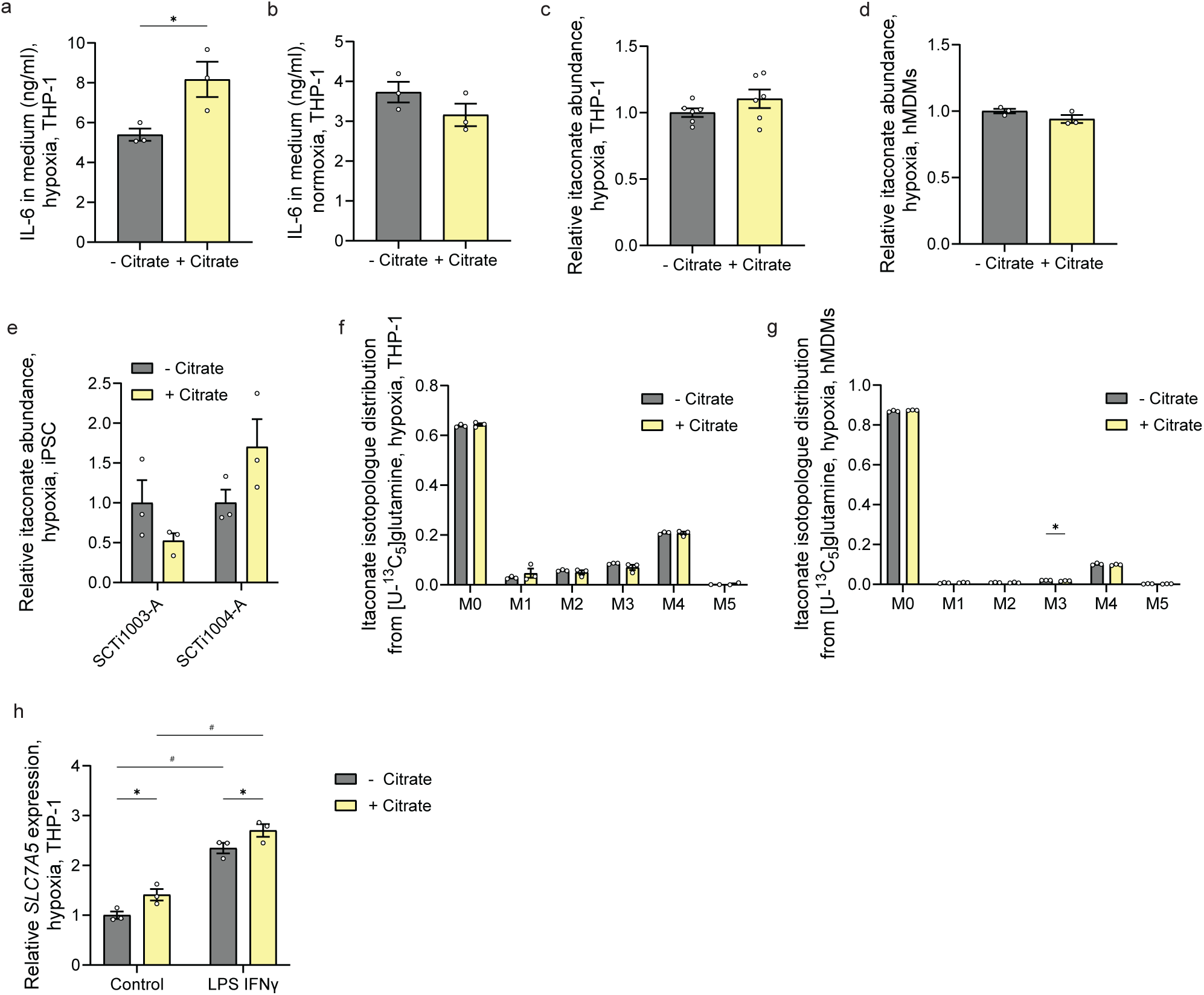
Extracellular citrate modulates inflammatory response. **a**, IL-6 concentration in media (ng/ml) of macrophage-like THP-1 cells. **b**, IL-6 concentration in media (ng/ml) of hMDMs. **c-e,** Relative intracellular abundance of itaconate in **c,** macrophage-like THP-1 cells, **d,** hMDMs, and **e,** iPSC-derived macrophages (SCTi1003-A, SCTi1004-A). **f,g**, Isotopologue distribution on itaconate from [U-^13^C_5_]glutamine in **f,** macrophage-like THP-1 cells and **g,** hMDMs. **h**, Expression of *SLC7A5* in macrophage-like THP-1 cells in hypoxic conditions. Cells were treated with 1 mM citrate and exposed to 100 ng/ml LPS and 400 U/ml IFNγ for 24 hours. Hypoxic conditions were performed with 2 % oxygen. Experiments were repeated independently three or more times with three (a,b,d-f) or six (c) cellular replicates each. Data are presented as means ± s.e.m. (a,b,d-f) or as box (25^th^ to 75^th^ percentile with median line) and whiskers (min. to max. values) (c). Statistical significances were determined with uncorrected Fisher’s LSD test with * *p* < 0.05; ** *p* < 0.01, *** *p* < 0.001, ^#^ *p* < 0.0001.

To further assess inflammatory metabolism, we quantified levels of the immunomodulatory metabolite itaconate, which is produced under pro-inflammatory conditions in macrophages. Immune-responsive gene 1 (IRG1) protein catalyzes the decarboxylation of cis-aconitate to itaconate. Since citrate supplies carbons to the cis-aconitate pool, citrate availability is directly linked to itaconate metabolism^30^. We observed that extracellular citrate did not affect itaconate abundances in macrophage-like THP-1 cells, hMDMs, and iPSC macrophages (Fig. 3c-e). Also, the labeling from [U-^13^C_5_]glutamine on itaconate was not affected by the addition of citrate (Fig. 3f, g). Consistently, ^13^C citrate tracing did not result in detectable labeling on itaconate (Fig. 2b), supporting that citrate is not used as a carbon source for TCA cycle metabolism and itaconate synthesis. Collectively, these data demonstrate that extracellular citrate enhances inflammatory cytokine production under hypoxic conditions.

To further characterize the impact of extracellular citrate on macrophage immunometabolism, we assessed levels of *SLC7A5* (large neutral amino acid transporter 1, LAT1). This transporter facilitates the exchange of intracellular glutamine for extracellular essential amino acids such as leucine and is associated with pro-inflammatory macrophage polarization, glutamine metabolism, and metabolic reprogramming^31^. We observed increased *SLC7A5* expression upon citrate supplementation under hypoxic conditions, which was further increased under inflammatory stimuli (Fig. 3h). These data are consistent with enhanced inflammatory activation and altered glutamine metabolism in human macrophages cultured in the presence of citrate.

Finally, to evaluate broader immune effects, we additionally examined T-cell activation markers. CD62L+ was significantly decreased with 5 mM citrate in unstimulated as well as stimulated T-cells (Sup. Fig. 3a). While TNFα was not affected in unstimulated T-cells, it was also significantly decreased with 5 mM citrate in stimulated T-cells. However, 1 mM citrate led to an increase of TNFα (Sup. Fig. 3b). Together, these data indicate that lower citrate doses promote pro-inflammatory-macrophage activation without acting as a direct metabolic substrate. However, higher doses of 5 mM may inhibit glycolysis, thereby affecting cytokine production. Further, our data indicate that extracellular citrate acts as a metabolic regulator of macrophages and links glutamine metabolism to functional immune responses.

### Extracellular citrate promotes intracellular growth of *S.* Typhimurium in co-culture with macrophages under hypoxic conditions

Given that extracellular citrate modulates macrophage metabolism and inflammatory responses, we next investigated whether these effects influence host-pathogen interactions. To investigate the functional consequences of extracellular citrate on bacterial infection and host-pathogen interaction, we established a co-culture system and infected macrophage-like THP-1 cells with *S.* Typhimurium (Fig. 4a). Since *S.* Typhimurium can use citrate as a carbon source^32–34^, we first confirmed that extracellular citrate does not directly enhance bacterial growth (Sup. Fig. 4a). Next, we assessed the effects of citrate in bacterial infection (Fig. 4a). Under normoxic conditions, *S.* Typhimurium infection decreased intracellular citrate abundance which was restored by extracellular citrate supplementation (Fig. 4b). As expected, hypoxia decreased citrate levels compared to normoxia. Further, infection in hypoxia did not further affect intracellular citrate levels and citrate supplementation did not significantly increase levels in infected cells (Fig. 4b). These data suggest that infection and hypoxia lead to similar effects on citrate metabolism.

**Figure 4:**
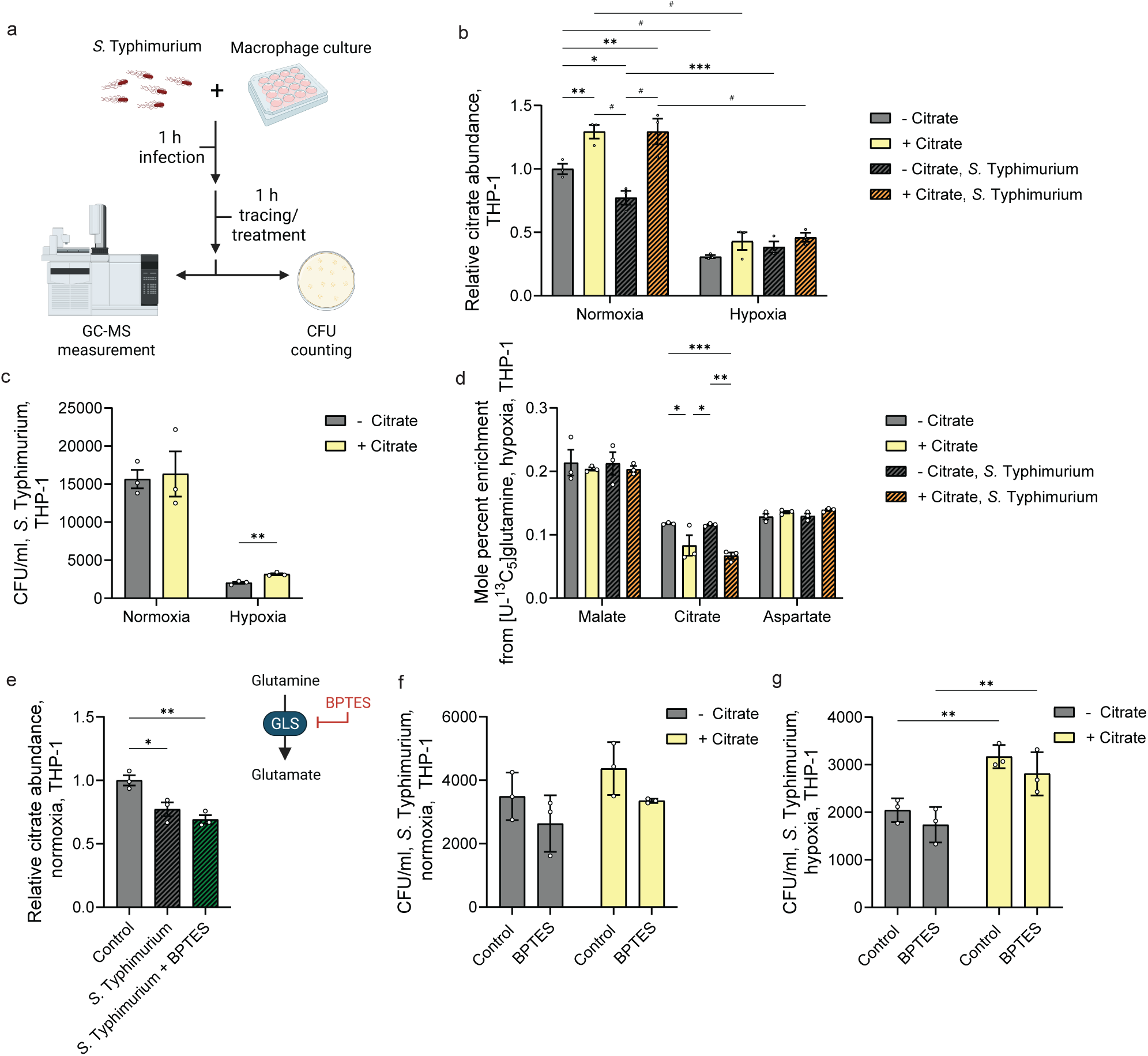
Extracellular citrate promotes intracellular growth of *S.* Typhimurium in co-culture with macrophages in hypoxia. **a,** Schematic illustration of infection experiments with *S.* Typhimurium in macrophage-like THP-1 cells. **b**, Relative intracellular citrate abundance in macrophage-like THP-1 cells in hypoxic and normoxic conditions after infection with *S.* Typhimurium. **c,** CFU/ml of *S.* Typhimurium after infection of macrophage-like THP-1 cells in hypoxic and normoxic conditions. **d,** Mole percent enrichment from [U-^13^C_5_]glutamine in macrophage-like THP-1 cells after infection with *S.* Typhimurium. **e,** Relative intracellular abundance of citrate with *S.* Typhimurium infection and 10 µM BPTES. **f,g,** CFU of *S.* Typhimurium after infection of macrophage-like THP-1 cells with extracellular citrate and 10 µM BPTES in **f,** normoxia and **g,** hypoxia. Cells were treated with 1 mM citrate and infected with MOI of 1:1 for 2 h. Experiments were repeated independently three or more times (b,c) or done once (d) or twice (e-g) with three cellular replicates each. Data are presented as means ± s.e.m.. Statistical significances were determined with uncorrected Fisher’s LSD test (c,f-g) or one-way (e) or two-way *ANOVA* (b,d) with * *p* < 0.05; ** *p* < 0.01, *** *p* < 0.001, ^#^ *p* < 0.0001.

To determine the functional impact of extracellular citrate on *S.* Typhimurium growth, we quantified intracellular bacterial load by colony-forming unit (CFU) after infection of macrophages. Under normoxic conditions, citrate had minor effects on intracellular bacterial load. Hypoxia significantly decreased CFU compared to normoxia, whereas citrate supplementation partially rescued bacterial growth under hypoxic conditions (Fig. 4c). Our findings are consistent with enhanced extracellular citrate uptake, as indicated by increased intracellular citrate levels, and function under hypoxia (Fig. 1, 2, 3). Given that extracellular citrate modulates macrophage metabolism (Fig. 2), we next asked whether these effects are associated with altered substrate utilization during infection.

To address this, we quantified metabolic flux and carbon utilization in our infection model, focusing on glutamine metabolism using [U-^13^C_5_]glutamine. We observed that extracellular citrate had minor effects on malate and aspartate labeling, indicating limited effects on glutamine utilization under these conditions (Fig. 4d). This observation contrasts our results obtained with LPS/IFNγ treated cultures (Fig. 2). This discrepancy may reflect the shorter tracing time in infection experiments (1 h trace) compared to inflammatory stimulation with LPS/IFNγ (24 h trace) due to the short doubling time of ∼20 min at 37 °C of *S.* Typhimurium^35,36^.

To test the potential contribution of glutamine metabolism to the observed phenotype, we inhibited glutamine utilization. Since our findings revealed that extracellular citrate reduced glutamine contribution to TCA cycle metabolism under inflammatory conditions (Fig. 2), we aimed to phenocopy this effect using the pharmacological glutaminase (GLS) inhibitor bis-2-(5-phenylacetamido-1,2,4-thiadiazol-2-yl)ethyl sulfide (BPTES). Inhibition of GLS reduced ^13^C incorporation of [U-^13^C_5_]glutamine into TCA cycle intermediates and itaconate, confirming effective inhibition of GLS activity (Sup. Fig. 4b). *S.* Typhimurium infection decreased intracellular citrate levels, which were further decreased with BPTES addition (Fig. 4e). Of note, BPTES did not affect bacterial growth (Sup. Fig. 4c). Further, BPTES had minor effects on intracellular bacterial load under either normoxic or hypoxic conditions, with or without citrate supplementation (Fig. 4f, g).

These data demonstrate that inhibition of GLS, resulting in decreased glutamine usage for TCA cycle metabolism, does not recapitulate the pro-bacterial effect of extracellular citrate. These findings suggesting that citrate-induced metabolic changes may create a permissive environment for bacterial survival under hypoxic conditions through mechanisms that are not solely explained by altered glutamine flux. This suggests a complex interplay between host metabolism and pathogen survival.

### Extracellular citrate modulates host-pathogen interactions during L. pneumophila infection

To determine whether these observed effects are conserved across intracellular pathogens, we next investigated host-pathogen interactions using *L. pneumophila*. *L. pneumophila* is a slower-growing bacterium compared to *S.* Typhimurium and enables metabolic tracing over almost 24 h with a doubling time of approximately 2 hours^36,37^ (Fig. 5a). We observed minor effects on bacterial growth in minimal medium supplemented with citrate, indicating that citrate does not directly promote bacterial proliferation (Sup. Fig. 5a). Instead, bacterial growth might be influenced by host metabolism and host-pathogen interactions.

**Figure 5:**
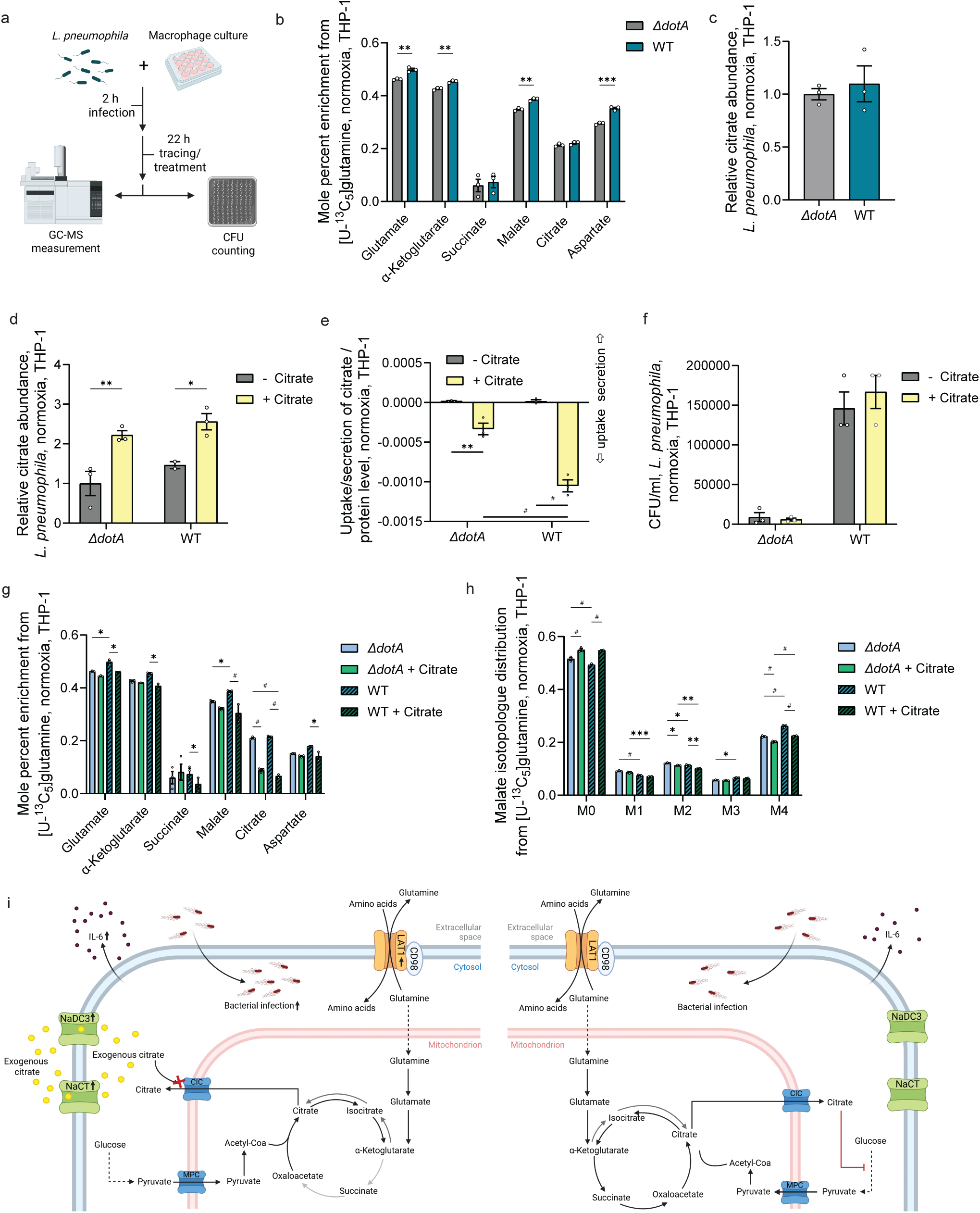
Extracellular citrate influences host-pathogen interactions in *L. pneumophila* infection models. **a**, Schematic illustration of infection experiments with *L. pneumophila* in macrophage-like THP-1 cells. **b**, Mole percent enrichment from [U-^13^C_5_]glutamine with *L. pneumophila ΔdotA* and WT infection. **c,** Relative citrate abundance with *L. pneumophila ΔdotA* and WT infection. **d,** Relative citrate abundance after *L. pneumophila ΔdotA* and WT infection with and without extracellular citrate. **e,** Citrate uptake after *L. pneumophila ΔdotA* and WT infection with and without extracellular citrate. **f,** CFU/ml of *L. pneumophila ΔdotA* and WT with and without extracellular citrate. **g,** Mole percent enrichment from [U-^13^C_5_]glutamine after infection with *L. pneumophila ΔdotA* and WT infection with and without extracellular citrate. **h,** Malate isotopologue distribution from [U-^13^C_5_]glutamine infection with *L. pneumophila ΔdotA* and WT infection with and without extracellular citrate. **i,** Schematic depicting impact of extracellular (left) and intracellular (right) citrate on macrophage metabolism and function. Experiments were independently repeated once (b, g-h), twice (c, d) or three times (e, f), with three cellular replicates each. Cells were treated with 1 mM citrate and infected with MOI of 1:10 for 22 h. Data are presented as means ± s.e.m.. Statistical significances were determined with uncorrected Fisher’s LSD test (b, c, f) or two-way *ANOVA* (e, g, h) with * *p* = 0.05; ** *p* < 0.01, *** *p* < 0.001, ^#^ *p* < 0.0001.

Similar to hypoxic conditions, bacterial infection has been associated with increased import and utilization of glutamine as a carbon source for TCA cycle metabolism, including succinate synthesis^38–40^. To isolate infection-driven metabolic effects, we established a co-culture system with *L. pneumophila* wild type (WT) and performed experiments under normoxic conditions. We compared our results to experiments with the non-infectious *ΔdotA* strain, which served as a control^41^. Our tracing experiments revealed that *L. pneumophila* (WT) infections increased label from ^13^C glutamine on TCA cycle intermediates and related amino acids (Fig. 5b). However, intracellular citrate abundance was not significantly affected by infection alone (Fig. 5c). Supplementation of extracellular citrate increased intracellular citrate levels with no difference between *ΔdotA* and WT infection (Fig. 5d). Notably, citrate uptake was significantly increased in WT-infected cells indicating that infection promotes citrate import (Fig. 5e). We also quantified bacterial load which was not affected by extracellular citrate supplementation (Fig. 5f).

Next, we applied a ^13^C glutamine tracer to investigate the impact of extracellular citrate on glutamine-dependent metabolism during infections with *L. pneumophila*. We observed that the labeling on TCA cycle intermediates and related amino acids was decreased with extracellular citrate in WT-infected cells. Labeling in *ΔdotA*-infected cells was less affected by extracellular citrate (Fig. 5g). Further, we observed that M4 labeling on malate was preferentially reduced by extracellular citrate (Fig. 5h), indicating that the oxidative TCA cycle metabolism (M4 on malate) is more affected than the reductive (M3 on malate). The contribution of ^13^C glucose-derived carbons into the TCA cycle was overall decreased with *L. pneumophila* infection, but extracellular citrate had minor effects (Sup. Fig. 5b).

In conclusion, we demonstrate that human macrophages import and accumulate extracellular citrate affecting immune responses and function. Further, extracellular citrate alters glutamine metabolism under conditions such as hypoxia and pro-inflammatory stimulation, which may influence intracellular bacterial growth and highlight a context-dependent role of citrate in host metabolic responses to intracellular pathogens (Fig. 5i).

## Discussion

Our results demonstrate that extracellular citrate is taken up by human macrophages and functions as a metabolic regulator rather than a major carbon source. Citrate rewires glutamine-dependent metabolism under stress conditions, modulates inflammatory responses, and exerts context-dependent effects on intracellular bacterial growth. These findings reveal a decoupling of nutrient uptake from metabolic utilization and demonstrate how extracellular metabolites influence immune cell metabolism.

Citrate and its transporters NaCT (encoded by *SLC13A5*) and NaDC3 (encoded by *SLC13A3*) have been implicated in metabolic and neurological diseases, including citrate transporter disease^15–18,42^. Extracellular citrate primarily affects tissues with high expression of citrate transporters, such as liver and brain. Notably, most cell culture media lack citrate despite its abundance in plasma^8^. Our data demonstrate that human macrophages accumulate extracellular citrate across multiple models including primary hMDMs and iPSC-derived macrophages. Under hypoxic conditions, increased citrate uptake together with elevated intracellular citrate levels and *SLC13A5* expression suggest that citrate may promote its own uptake in a context-dependent manner.

In hepatocytes, citrate is used as a carbon source for TCA cycle metabolism and lipogenesis under hypoxic conditions^8^. In contrast, our study demonstrates that extracellular citrate uptake is uncoupled from its utilization as a carbon source in human macrophages. Instead, extracellular citrate altered metabolic fluxes, indicating that small molecules may also act as regulators of metabolic networks rather than as sole substrates. Mechanistically, extracellular citrate reduced glutamine metabolism and utilization under hypoxic and inflammatory conditions. Since glutamine is a major substrate in activated macrophages, citrate may modulate central carbon metabolism by limiting glutamine utilization^4,38–40,43,44^. This is further supported by increased expression of *SLC7A5* in the presence of citrate, as this transporter modulates glutamine exchange and is linked to metabolic reprogramming and inflammation in macrophages^31,45^. These findings suggest that extracellular citrate influences the homeostasis of nutrient usage for mitochondrial metabolism. Further, they indicate that metabolic regulation can occur independently of substrate utilization, highlighting the importance of isotope-labeled studies in immunometabolism^46^.

Previous studies reported that extracellular citrate promotes the secretion of cytokines including IL-1β, IL-10, and TNFα^16–18,42^. In our human macrophage experiments, extracellular citrate enhanced IL-6 cytokine release while promoting intracellular growth of *S.* Typhimurium under hypoxic conditions. This suggests that increased inflammatory activation does not necessarily translate into improved antimicrobial defense. Instead, metabolic rewiring induced by citrate may create a cellular environment that is permissive for bacterial survival. Citrate had less impact on *L. pneumophila* replication, potentially due to citrate usage for lipid synthesis and vacuole formation^47^. These findings demonstrate the complex interplay between host metabolism and pathogen survival and suggest that metabolic regulation has divergent functional outcomes depending on the infectious context.

Since macrophages respond to extracellular citrate by altering metabolic flux and immune function, elevated circulating citrate levels, as observed in SLC13A5 citrate transporter disorder^3,12^, may influence immune responses in vivo. More broadly, our data suggest that extracellular metabolites such as citrate can act as systemic regulators of immune cell metabolism, linking metabolic disorders to immune function through altered substrate availability. Further work is needed to define transporter contributions under different conditions. NaDC3 transports itaconate^48,49^, suggesting that SLC13 activity may coordinate citrate uptake with immunometabolic regulation and inflammatory signaling.

In conclusion, our study identifies extracellular citrate as a previously underappreciated regulator of macrophage metabolism. By decoupling uptake from metabolic utilization, citrate modulates glutamine utilization, immune responses, and host-pathogen interactions. Together, these findings challenge the view that extracellular nutrients primarily serve as metabolic substrates and instead highlight extracellular metabolites as context-dependent regulators of immune cell metabolism and function. This concept expands current understanding of immunometabolism and may inform therapeutic strategies targeting citrate metabolism.

## Ethics statement

Blood samples were obtained from healthy donor volunteers under the Research Ethics Committee permission (HR/DP-21/22–14568). All subjects gave their written informed consent. The study was conducted in accordance with the Declaration of Helsinki. All storage of samples obtained complied with the requirements of the Data Protection Act 1998 and the Human Tissue Act 2004, issued by the UK parliament.

## Acknowledgments

We thank all members of the Cordes Lab for helpful discussions. Figures were created with BioRender.com. This study was supported, in part, by the Deutsche Forschungsgemeinschaft (DFG, German Research Foundation, Project Number 555979574) (to T.C.), the German Federal Ministry for Economic Affairs and Climate Action (BMWK, ZIM grant 16KN098645) (to M.S.), and the German Federal Ministry of Research, Technology and Space (BMFTR) under grant number 01KX2324 as part of the project ‘Microbial Stargazing’ (to J.P.). We acknowledge support from the Open Access Publication Funds of Technische Universität Braunschweig.

## Availability of data

All data associated with the study are in the manuscript. Source data are deposited in the repository platform of Technische Universität Braunschweig with a dedicated DOI upon acceptance of the manuscript.

## Declaration of Generative AI and AI-assisted technologies in the writing process

During the preparation of this work, the authors used a generative AI tool (ChatGPT) to assist with language and readability. The authors reviewed and edited the content as needed and take responsibility for the content of the publication.

## Author contributions

H.F.V. – Conceptualization, Methodology, Investigation, Formal Analysis, Visualization, Writing – Original Draft, Writing – Review & Editing; F.L. – Investigation and Formal Analysis related to *L. pneumophila* experiments; S.W. – Investigation; S.N. – Investigation and Formal Analysis related to T-cells experiments; H.G. – Resources provided hPBMCs; K.O.A., K.M., J.P. – Resources provided iPSC lines; R.V.T. – Methodology, Formal Analysis and Resources related to untargeted metabolomics analysis; A.S. – Methodology and Resources related to T-cell experiments; M.S. – Methodology and Resources related to *L. pneumophila* experiments; T.C. – Conceptualization, Methodology, Investigation, Formal Analysis, Visualization, Writing – Original Draft, Writing – Review & Editing, Funding Acquisition, Resources, Funding Acquisition, Supervision; All authors have read and agreed to the published version of the manuscript.

## List of Abbreviations

[2,4-^13^C_2_]citrate: 2,4 labeled citrate
[U-^13^C_5_]glutamine: Uniformly labeled glutamine
[U-^13^C_6_]glucose: Uniformly labeled glucose
^12^C: Carbon 12
^13^C: Carbon 13
ACES: N-(2-acetamido)-2-aminoethanesulfonic acid
ACLY: ATP-citrate-lyase
ATCC: American Type Culture Collection
BCA: Bicinchoninic Acid Assay
BCYE: Buffered charcoal yeast extract
BPTES: Bis-2-(5-phenylacetamido-1,2,4-thiadiazol-2-yl)ethyl sulfide
BSA: Bovine serum albumin
CFU: Colony-forming units
CoA: Coenzyme A
cRPMI: Complete Roswell Park Memorial Institute medium
CSF: Cerebrospinal fluid
CT: Cycling threshold
DAMP: Damage-associated molecular pattern
DMEM: Dulbecco’s Modified Eagle Medium
ELISA: Enzyme-linked immunosorbent assay
FAMEs: Fatty acid methyl esters
FBS: Fetal bovine serum
FCCP: Trifluoromethoxy carbonyl cyanide phenylhydrazone
GC: Gas chromatography
GLS: Glutaminase
hMDMs: Human monocyte-derived macrophages
HRP: Horse radish peroxidase
IFNγ: Interferon γ
IL-6: Interleukin 6
IPSC: Induced pluripotent stem cell
*L. pneumophila*: *Legionella pneumophila* Corby
LPS: Lipopolysaccharide
LRS: Leucocyte reduction system
MASLD: Metabolically-dysfunction-associated steatotic liver disease
MOI: Multiplicity of Infection
MS: Mass spectrometry
MTBSTFA: *N*-tert-Butyldimethylsilyl-*N*-methyltrifluoracetamide
NaCT: Sodium-dependent citrate transporter (SLC13A5)
NaDC1: Sodium-dependent dicarboxylate transporter 1 (SLC13A2)
NaDC3: Sodium-dependent dicarboxylate transporter 3 (SLC13A3)
NAFLD: Non-alcoholic fatty liver disease
OCR: Oxygen consumption rate
PMA: Phorbol 12-myristate 13-acetate
P/S: Penicillin/Streptomycin
PBMC: Peripheral Blood Mononuclear Cell
PBS: Phosphate-buffered saline
Rh M-CSF: Recombinant human macrophage colony-stimulating factor
RIPA: Radioimmunoprecipitation Assay buffer
RPMI: Roswell Park Memorial Institute medium
*RPL27*: Ribosomal protein L27 (human gene)
RT: Room temperature
s.e.m.: Standard error of mean
*S.* Typhimurium: *Salmonella enterica* Typhimurium
SDH: Succinate dehydrogenase
*SLC7A5*: Solute carrier family 7 member 5 (human gene)
*SLC13A3*: Solute carrier family 13 member 3 (human gene)
*SLC13A5*: Solute carrier family 13 member 5 (human gene)
tBDMS: Tert-Butyldimethylsilylchloride
TCA: Tricarboxylic acid
TNFα: Tumor necrosis factor alpha
UHPLC: Ultra-high performance liquid chromatography
WT: Wild type
YEB: Yeast extract beef

## Supplementary Figures

**Supplementary Figure 1:**
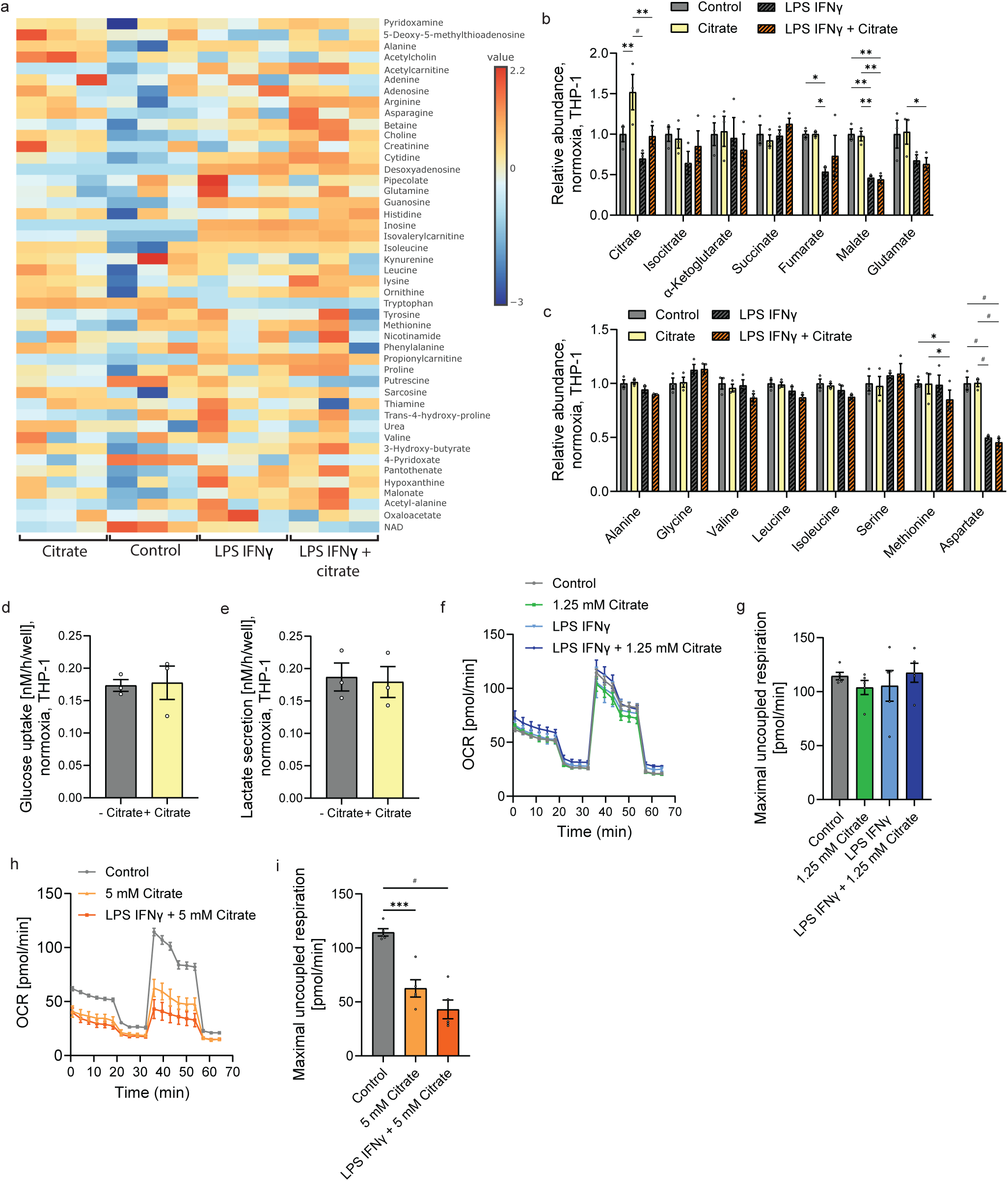
Extracellular citrate has minor effects on metabolism under normoxic conditions. **a**, Heatmap depicting fold change measured by untargeted LC-MS of metabolites in macrophage-like THP-1 cells with and without citrate and LPS/IFNγ. **b,c**, Relative intracellular abundance of **b**, TCA cycle intermediates, and **c**, amino acids in macrophage-like THP-1. **d,** Glucose uptake by macrophage-like THP-1. **e,** Lactate secretion by macrophage-like THP-1. **f,** Oxygen consumption rate of macrophage-like THP-1 cells with and without 1.25 mM citrate and LPS/IFNγ. **g,** Maximal uncoupled respiration of macrophage-like THP-1 cells with and without 1.25 mM citrate and LPS/IFNγ. **h,** Oxygen consumption rate of macrophage-like THP-1 cells with and without 5 mM citrate and LPS/IFNγ. **i,** Maximal uncoupled respiration of macrophage-like THP-1 cells with and without 5 mM citrate and LPS/IFNγ. Cells were treated with 1 mM citrate and exposed to 100 ng/ml LPS and 400 U/ml IFNγ for 24 hours. Data baseline corrected to control without LPS/IFNγ and without citrate (b-c). Experiments were independently repeated one (a), three (d-i) or more times (b, c) with three cellular replicates. Data are presented as means ± s.e.m.. Statistical significances were determined with two-way *ANOVA* (b, c) or uncorrected Fisher’s LSD test (c, e) with * *p* < 0.05; ** *p* < 0.01, *** *p* < 0.001, ^#^ *p* < 0.0001.

**Supplementary Figure 2:**
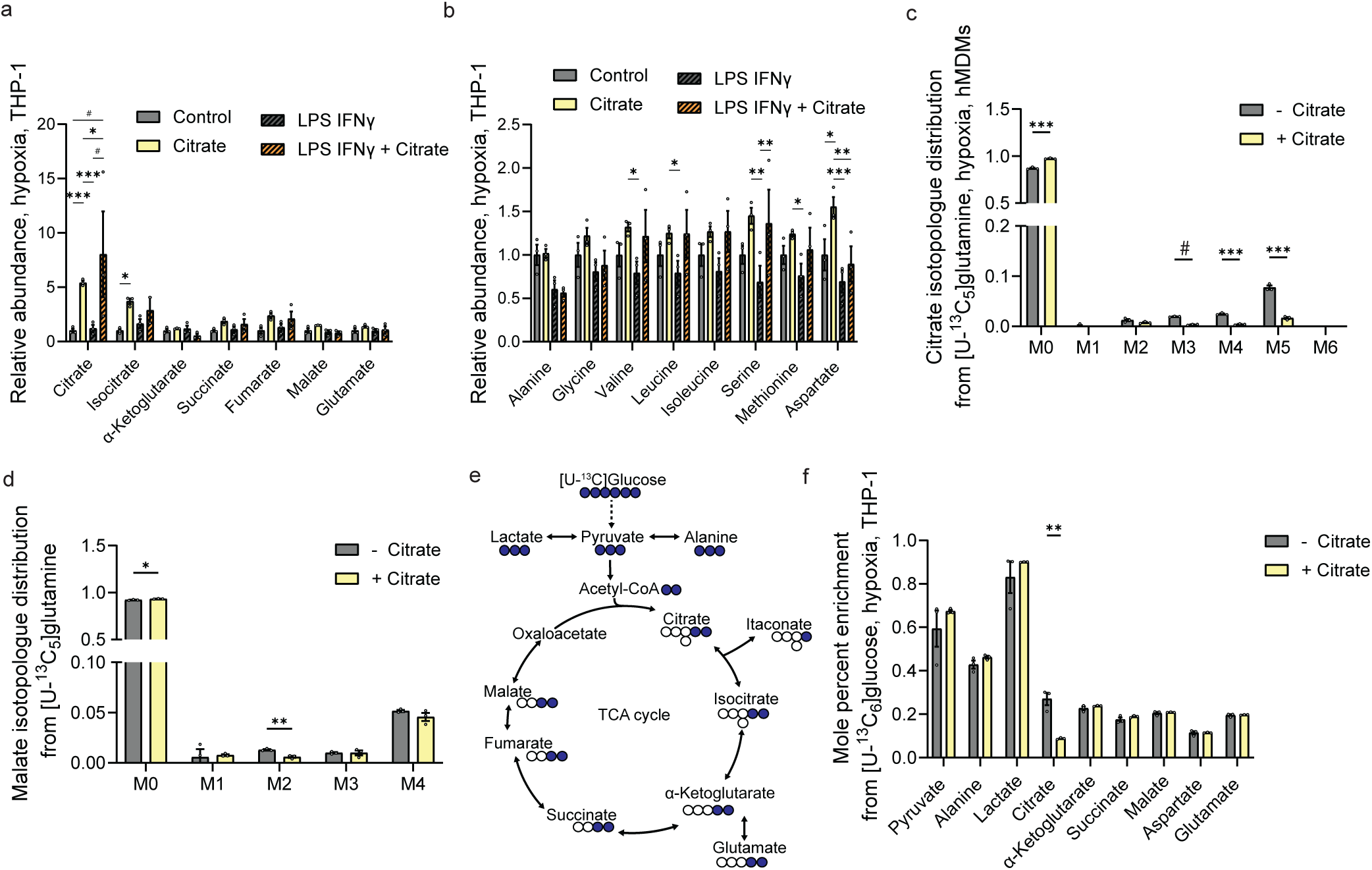
Citrate uptake influences central carbon metabolism. **a,b**, Relative intracellular abundance of **a**, TCA cycle intermediates and **b**, amino acids in macrophage-like THP-1 cells. **c**, Citrate isotopologue distribution from [U-^13^C_5_]glutamine in hMDMs in hypoxia. **d,** Malate isotopologue distribution from [U-^13^C_5_]glutamine in hMDMs. **e,** Schematic depicting the use of [U-^13^C_6_]glucose as carbons for TCA cycle intermediates. White circles: ^12^C; blue circles: ^13^C. **f,** Mole percent enrichment from [U-^13^C_6_]glucose on TCA cycle intermediates in macrophage-like THP-1 cells. Experiments were independently repeated two (a, b) or more times (d) with three cellular replicates. Cells were treated with 1 mM citrate and exposed to 100 ng/ml LPS and 400 U/ml IFNγ for 24 hours. Data are presented as means ± s.e.m.. Statistical significances were determined with uncorrected Fisher’s LSD test with * *p* < 0.05; ** *p* < 0.01, *** *p* < 0.001, ^#^ *p* < 0.0001.

**Supplementary Figure 3:**
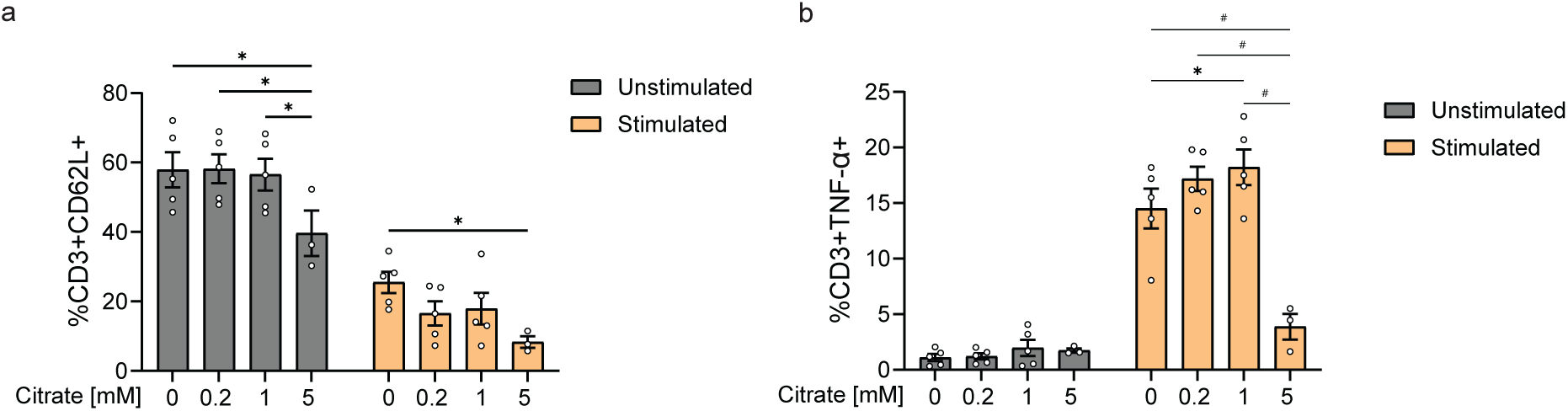
High citrate levels influence T-cell immune response. **a**, CD3+CD69L^+^ and **b**, CD3+TNFα^+^ in PBMCs treated with 0 mM, 0.2 mM, 1 mM, and 5 mM citrate. Experiments were independently repeated with 5 donors. Data are presented as means ± s.e.m.. Statistical significances were determined with two-way *ANOVA* with * *p* < 0.05; ** *p* < 0.01, *** *p* < 0.001, ^#^ *p* < 0.0001.

**Supplementary Figure 4:**
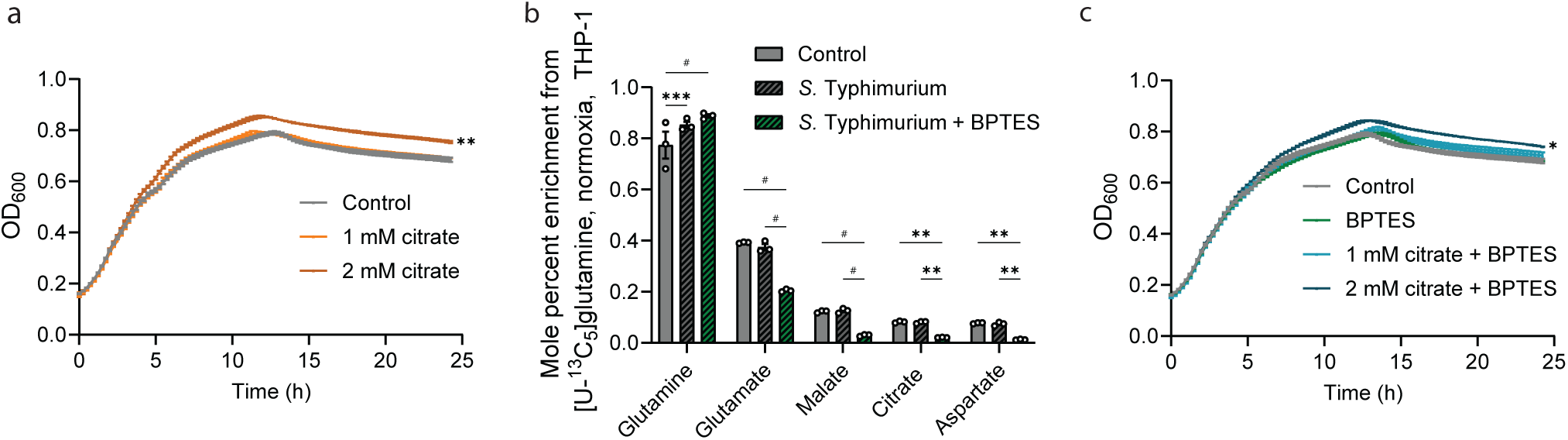
Impact of citrate and BPTES on growth and metabolism of *S.* Typhimurium. **a,** Growth curve of *S.* Typhimurium with 1 mM and 2 mM citrate in RPMI medium. **b,** Mole percent enrichment from [U-^13^C_5_]glutamine with *S.* Typhimurium infection and 10 µM BPTES. **e**, Growth curve of *S.* Typhimurium with 1 mM and 2 mM citrate and 10 µM BPTES in RPMI medium. Experiments were independently repeated one (b) or three times (a,c) with three (b) cellular or five technical replicates (a,c). Cells were treated with 1 mM citrate and infected with MOI of 1:1 for 2 h. Data are presented as means ± s.e.m.. Statistical significances were determined with two-way *ANOVA* with * *p* < 0.05; ** *p* < 0.01, *** *p* < 0.001, ^#^ *p* < 0.0001.

**Supplementary Figure 5:**
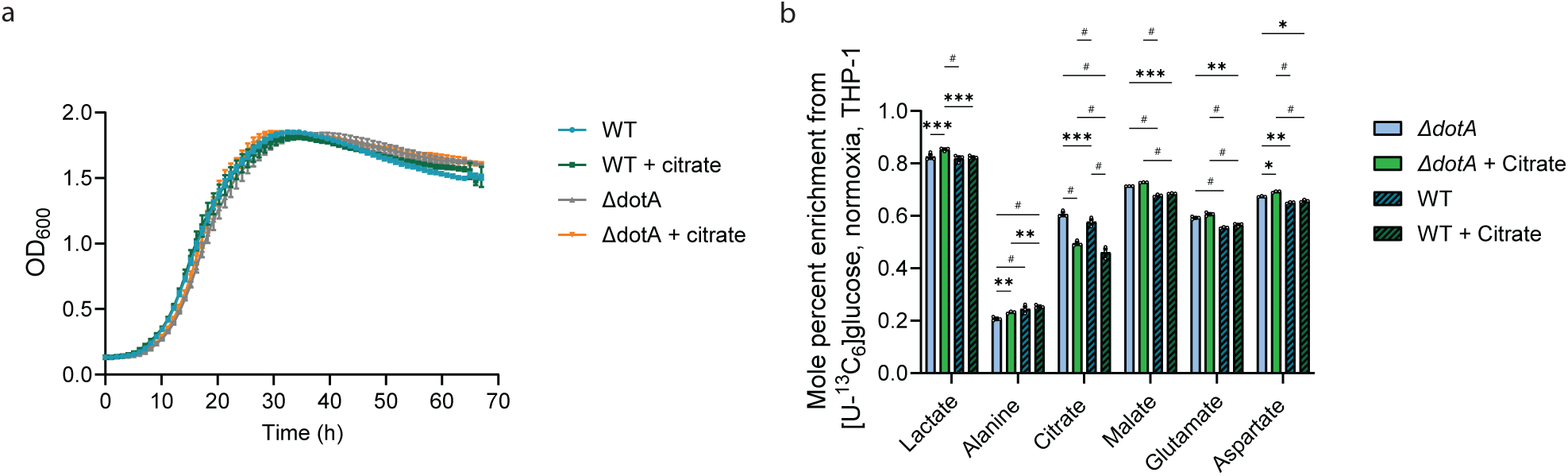
Effects of extracellular citrate on *L. pneumophila*. **a**, Growth curve of *L. pneumophila ΔdotA* and wild type (WT) in minimal medium. **b,** Mole percent enrichment from [U-^13^C_6_]glucose with *L. pneumophila ΔdotA* and WT infection. Experiments were repeated independently two (b) or three times (a), with five technical replicates (a) or three cellular replicates (b). Cells were treated with 1 mM citrate and infected with MOI of 1:10 for 22 h. Data are presented as means ± s.e.m.. Statistical significances were determined with two-way *ANOVA* (b) with * *p* < 0.05; ** *p* < 0.01, *** *p* < 0.001, ^#^ *p* < 0.0001.

